# Elevated blood pressure in high-fat diet-exposed low birthweight rat offspring is most likely caused by elevated glucocorticoid levels

**DOI:** 10.1101/2020.02.21.958884

**Authors:** Takahiro Nemoto, Takashi Nakakura, Yoshihiko Kakinuma

## Abstract

Being delivered as a low birthweight (LBW) infant is a risk factor for elevated blood pressure and future problems with cardiovascular and cerebellar diseases. Premature babies have been reported to possess a lower number of nephrons, but the mechanisms by which blood pressure is elevated in full-term LBW infants remain unclear. We generated a fetal low-carbohydrate and calorie-restricted model rat, and some individuals showed postnatal growth failure caused by increased miR-322 expression in the liver and decreased growth hormone receptor expression. Using this model, we examined how a high-fat diet-induced mismatch between prenatal and postnatal environments could elevate blood pressure after growth. Although LBW rats fed standard chow had slightly higher blood pressure than control rats, their blood pressure was significantly higher than controls when exposed to a high-fat diet. Observation of glomeruli subjected to PAM staining showed no difference in number or size. Aortic and cardiac angiotensin II receptor expression was altered with compensatory responses. Blood aldosterone levels were not different between control and LBW rats, but blood corticosterone levels were significantly higher in the latter with high-fat diet exposure. Administration of metyrapone, a steroid synthesis inhibitor, reduced blood pressure to levels comparable to controls. We showed that high-fat diet exposure causes impairment of the pituitary glucocorticoid feedback via miR-449a. These results clarify that LBW rats have increased blood pressure due to high glucocorticoid levels when they are exposed to a high-fat diet. These findings suggest a new therapeutic target for hypertension of LBW individuals.

## Introduction

Hypertension is a multifactorial disease caused by the interaction of genes and environment, and is a major factor in the future development of cardiovascular disease (CVD)^1, 2^. Risk factors for developing hypertension vary, but one is considered to be low birthweight (LBW). A systematic review by Huxley *et al.* found that birthweight and blood pressure are negatively correlated^3^. While studying the morbidity and mortality of a large cohort in the general population, Barker and colleagues found a strong association between birthweight and susceptibility to both adult onset hypertension and CVD^4^. Many other epidemiological and experimental studies have strongly supported the Barker hypothesis^5–8^. Furthermore, the developmental origins of health and disease (DOHaD) theory has evolved into the idea that mismatches between the acquired constitution and the growth environment create a risk of developing noncommunicable diseases^9–13^.

In Japan, an increase in the rate of LBW infants has been reported, and it is an urgent task to control the future development of non-communicable chronic diseases in these children^14–16^. However, verification of the mechanisms underlying the adult onset of hypertension caused by LBW in humans is time consuming and has many ethical problems. Therefore, we generated a model rat that was fed a low-carbohydrate and calorie-restricted diet during pregnancy. Although the rats did not deliver prematurely, the offspring weighed less and some of them failed to catch up during postnatal growth. In addition, we found that when they were exposed to restraint stress, blood corticosterone levels were further elevated due to a dysfunctional glucocorticoid-mediated feedback system in the pituitary gland^17, 18^. Therefore, the purpose of the present study was to clarify whether LBW caused by low carbohydrate and calorie restriction in fetuses increases blood pressure after growth, and if a mismatch between the acquired constitution and the growth environment further increases blood pressure, and which organs are involved.

The mechanisms that mediate fetal programming of hypertension have been extensively studied in three major organs or systems^19–21^: the kidney (decreased nephron number, activation of the renin-angiotensin system, or increased renal sympathetic nerve activity), the vasculature (alterations in structure or impaired vasodilation), and the neuroendocrine system (hypothalamic-pituitary-adrenal upregulation or altered adaptation to stress). Adverse events that occur in the uterus can affect the development of the fetal kidney and reduce the final number of nephrons^22^. Births of a low weight or that are premature are associated with an increased risk of hypertension, proteinuria, and kidney diseases that develop later in life. Therefore, we investigated the morphology of the kidneys of LBW rats and examined the number and size of glomeruli. We also investigated the expression levels of the angiotensin II receptors (AT1 and AT2), which are blood pressure regulators, in the aorta and heart. AT1 and AT2 are both G protein-coupled receptors, but AT2 shares only 34% identity with AT1^23, 24^. AT2 expression is found in the kidney and nervous and cardiovascular systems of adult rats^25, 26^. AT2 inhibits the AT1-mediated increase in inositol triphosphate and shows vasodilator, antifibrotic, antiproliferative, and anti-inflammatory effects^27–30^. Overexpression of AT2 promotes cardiac repair after myocardial infarction in mice^31^, and increased AT2 expression is thought to play an important role in tissue repair and regeneration as observed in response to injury and pathophysiological remodeling^32–36^.

In this study, we also performed immunostaining of arterial smooth muscle using an anti-α-smooth muscle (SMA) antibody. Well-established animal models indicate that environmentally induced intrauterine growth restriction (IUGR; by diet, diabetes, hormonal exposure, or hypoxia) increases the risk of development of various diseases in target organs later in life^37^. This suggests that a series of perturbations in fetal and postnatal growth may influence developmental programming and produce abnormal phenotypes. The endothelium, composed of endothelial cells, may be an important player for modulating long-term remodeling and the elastic properties of arterial walls. According to some clinical and experimental studies, fetuses, neonates, children, and adolescents who suffered from IUGR show signs of endothelial dysfunction (aortic intima-media thickness, carotid intima-media thickness and rigidity).

Finally, we examined the relationship between blood adrenal steroid levels and blood pressure in high-fat diet-exposed LBW rats. We previously reported that LBW rats maintain elevated blood corticosterone under restraint stress^17, 18^. Therefore, we investigated whether a high-fat diet, which is a chronic metabolic stress, could increase blood corticosterone levels in LBW rats. We have already shown that pituitary miR-449a is involved in the negative feedback of glucocorticoids^38^, and restraint stress-induced LBW rats have impaired induction of miR-449a expression in the anterior pituitary gland. Therefore, the expression of miR-449a in the anterior pituitary gland of high-fat diet-exposed LBW rats was examined. Finally, we investigated whether the administration of metyrapone, a steroid synthesis inhibitor, could normalize blood pressure.

## Methods

### Animals

Wistar rats were maintained at 23 ± 2 °C with a 12:12-h light-dark cycle (lights on at 0800 h, off at 2000 h). They were allowed *ad libitum* access to laboratory chow and distilled water. All experimental procedures were reviewed and approved by the Laboratory Animals Ethics Review Committee of Nippon Medical School. All experiments were performed in accordance with relevant guidelines and regulations^39^. We previously generated fetal low-carbohydrate and calorie-restricted rats^40^. Briefly, proestrus female rats (age, 9 weeks) were mated with normal male rats. Dams were housed individually with free access to water and were divided into two groups: low-carbohydrate and calorie-restricted diet (LC) dams were restricted in their calorie intake to 60% of the control group (D08021202, Research Diet Inc., New Brunswick, NJ), while control dams freely accessed food during the entire gestational period. The number of births was adjusted to 10~12 per mother rat, and the mother rats after birth were given standard chow by free-feeding in both groups. Rats were divided into two groups at 5 weeks of age, one on a high-fat diet (D12451, Research Diet) and the other on a standard diet. At 18 weeks of age, blood pressure was measured, blood was collected, organs were removed from the rats, and various analyses were performed.

### Tissue staining

The aortas and kidneys of rats were removed and fixed by immersion in 4% paraformaldehyde in 0.1 M phosphate buffer (pH 7.4) for 1 day at 4 °C, dehydrated through a graded ethanol series, and embedded in paraffin. The sections were cut with a microtome (SM 2000 R, Leica Biosystems, Wetzlar, Germany) and placed on PLATINUM PRO slides (Matsunami, Osaka, Japan) as previously described^41^. For observation of the renal glomerular basement membranes, kidney sections (1 μm thick) were deparaffinized and stained with periodic acid-methenamine silver (PAM). For immunohistochemistry, deparaffinized aorta sections were treated for antigen retrieval by heating in an autoclave in 1 mM EDTA at 121 °C for 5 min, and were then incubated overnight at 25 °C with mouse anti-α-smooth muscle (1:200; A5228, Sigma) in phosphate-buffered saline (PBS) containing 1% bovine serum albumin. After washing with PBS, the sections were incubated with Cy3-labeled donkey anti-rabbit IgG, Alexa Fluor 488-labeled donkey anti-mouse IgG (Jackson Immunoresearch, West Grove, PA, USA), and 4’,6-diamidino-2-phenylindole (DAPI; Dojindo, Kumamoto, Japan) for 2 h at room temperature. The specimens were examined with a BX53 microscope equipped with a DP80 microscope digital camera and cellSens imaging software (Olympus Optical, Tokyo, Japan).

### Blood pressure measurement

Blood pressure was measured non-invasively from tail blood volume, flow, and pressure using a volume pressure recording sensor and an occlusion tail cuff (CODA System; Hakubatec Lifescience Solutions, Tokyo, Japan)^42^. As reported previously, this is a highly accurate system that can non-invasively and simultaneously measure systolic and diastolic blood pressure and heart rate^43^. Prior to measurement, rats were placed on a 37 °C heating pad until the tail temperature reached 37 °C. After heating, blood pressure was measured 10 times, and the average value was used. All measurements were performed at the same time (10:00 am to 02:30 pm).

### RNA extraction and real-time RT-PCR

We performed mRNA and miRNA quantification as previously reported^17^. Total RNA was extracted from rat infrarenal aortas, hearts, kidneys, and pituitaries using RNAiso Plus (Takara, Shiga, Japan). The absorbance of each sample at 260 nm and 280 nm was assayed, and RNA purity was judged as the 260/280 nm ratio (The 260 /280 nm ratio of all samples used in this study was higher than 1.7). For miRNA expression analysis, first-strand cDNA was synthesized at 37 °C for 1 h using 1 μg of denatured total RNA and then terminated at 85 °C for 5 min using a Mir-X^®^ miRNA First-Strand Synthesis and SYBR^®^ qRT-PCR kit (Clontech Laboratories Inc., Mountain View, CA). For mRNA expression analyses, first-strand cDNA was generated using 0.5 μg of denatured total RNA; the reaction mixture was incubated at 37 °C for 15 min, 84 °C for 5 sec, and 4 °C for 5 min using a PrimeScript^®^ RT reagent kit with gDNA Eraser (Takara). PCR was performed by denaturation at 94 °C for 5 sec and annealing-extension at 60 °C for 30 sec for 40 cycles using SYBR premix Ex Taq (Takara) and specific primer sets for rat AT1 (RA061027, Takara), AT2 (RA060278, Takara), or GAPDH (RA015380, Takara). To normalize each sample for RNA content, GAPDH, a housekeeping gene, and U6 small nuclear RNA (Clontech Laboratories, Inc.) were used for mRNA and miRNA expression analyses, respectively. The 2^nd^ derivative method was used as the standard and for calculating C_t_ values, respectively^44^.

### Western blotting

We performed western blotting quantification as previously reported^40^. Protein samples from rat infrarenal aortas and hearts were extracted using complete lysis-M (Roche, Mannheim, Germany). The protein concentrations of lysate samples were determined using the Pierce 660 nm Protein assay (Thermo Scientific, Rockford, IL). Each 5 µg of protein was electrophoresed on a 5-20% gradient SuperSep^TM^ SDS-polyacrylamide gel (FUJIFILM Wako Pure Chemical Corporation, Osaka, Japan) and transferred to a nitrocellulose membrane. The transfer membranes were blocked with 5% skim milk and then incubated with an anti-AT1 antibody (1:1,000, GTX89149, GeneTex, Inc., Irvine, CA) or anti-AT2 antibody (1:1,000, GTX62361, GeneTex) for 1 h at room temperature. The transfer membranes were washed with TBS-T and further incubated with HRP-labeled anti-goat or anti-rabbit IgG (1:2,000, Jackson ImmunoResearch, West Grove, PA) for 1 h at room temperature. The signals were detected using SuperSignal West Dura extended duration substrate (Thermo Scientific). The membranes after detection were stripped of the antibody using Restore plus western blot stripping buffer (Thermo Scientific). Then, signals were detected again using THE^TM^ [HRP]-labeled β-actin antibody (1:1,000, A00730-40, GeneScript, Piscataway, NJ). The expression levels of AT1 or AT2 were quantified by correcting the AT1 or AT2 signal with the β-actin signal.

### Measurement of blood aldosterone and corticosterone levels

Aldosterone and corticosterone levels were measured using blood plasma from decapitated rats. Aldosterone was measured using a rat aldosterone EIA kit (#501090, Cayman Chemical, Ann Arbor, MI), and corticosterone was measured using a rat corticosterone ELISA kit (#501320, Cayman Chemical).

### Metyrapone administration

Metyrapone was purchased from Sigma-Aldrich (St Louis, MO). Administration of metyrapone to rats was performed according to a previous report^45^. Rats treated with metyrapone (10 mg/100 g body weight, s.c., twice daily at 0900 h and 1800 h for a week).

### Statistical analysis

Unpaired *t* tests or a one-way analysis of variance (ANOVA) followed by Turkey’s *post hoc* test for multiple comparisons were used for each statistical analysis. Prism 5.0 software (GraphPad Software, Inc., La Jolla, CA) was used for all calculations. Real-time RT-PCR and western blot data are expressed as percents ± SEM with the control set to 100. p <0.05 was considered statistically significant.

## Results

### Blood pressure

The blood pressure of rats fed the standard diet showed a slight but not significant increase in both systolic and diastolic pressure in LBW rats compared with control rats (Fig. 1A). When rats were exposed to a high-fat diet, the systolic and diastolic blood pressures of LBW rats were significantly higher than those of control rats (Fig. 1B).

**Figure 1.**
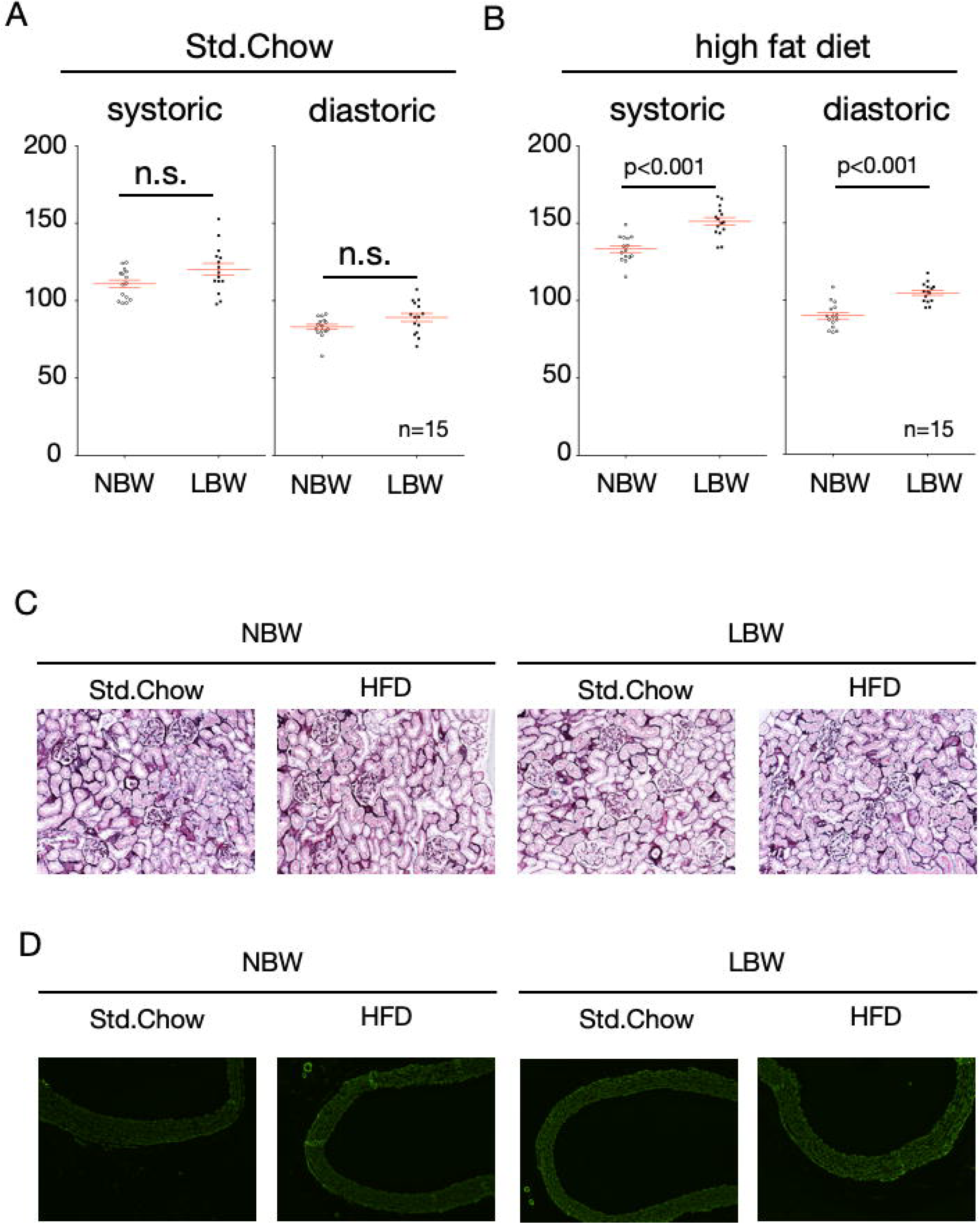
Blood pressure and tissue observations of kidneys and aortas of LBW rats with high-fat diet exposure. LBW rats were exposed to a high-fat diet (HFD) for 8 weeks. Blood pressure was measured by the tail-cuff method. (A) shows control rats (NBW) and LBW rats fed a standard chow, and (B) shows control rats and LBW rats exposed to a high-fat diet (n=15). Characteristic kidney PAM staining image and an aortic anti-SMA immunostaining image are shown (C and D).

### Renal glomerular morphology and aortic vascular smooth muscle staining

The kidneys of LBW rats and control rats exposed to a high-fat diet were excised and subjected to PAM staining. There was no difference in the number of glomeruli per field of view or in the size of the glomeruli, leading to no identification of glomerular hypertrophy, glomerular injury with mesangial proliferation, or matrix deposition (Fig. 1C). Similarly, aortic smooth muscle was stained with anti-SMA antibody, but no difference was found in smooth muscle thickness among them (Fig. 1D).

### Angiotensin II receptor expression

The expression levels of angiotensin II receptors (AT1 and AT2) in aortas and hearts were examined by real-time RT-PCR and western blotting. The expressions of mRNA and protein of AT1 in the aortas and hearts of LBW rats fed the standard diet were significantly lower than those in controls, respectively (Fig. 2A, 2C, 3A and 3C). When a high-fat diet was provided, their expressions were further decreased (Fig. 2A, 2C, 3A and 3C). While the expression of AT2 in the aortas of LBW rats fed a standard diet was lower than that in the controls (Fig. 2B and 2D), the expression of AT2 in the hearts of LBW rats fed a standard diet was higher than that in the controls (Fig. 3B and 3D). Exposure to a high-fat diet reduced their expression in the aorta (Fig. 2C and 2D), but significantly increased protein expression in the heart (Fig. 3D).

**Figure 2.**
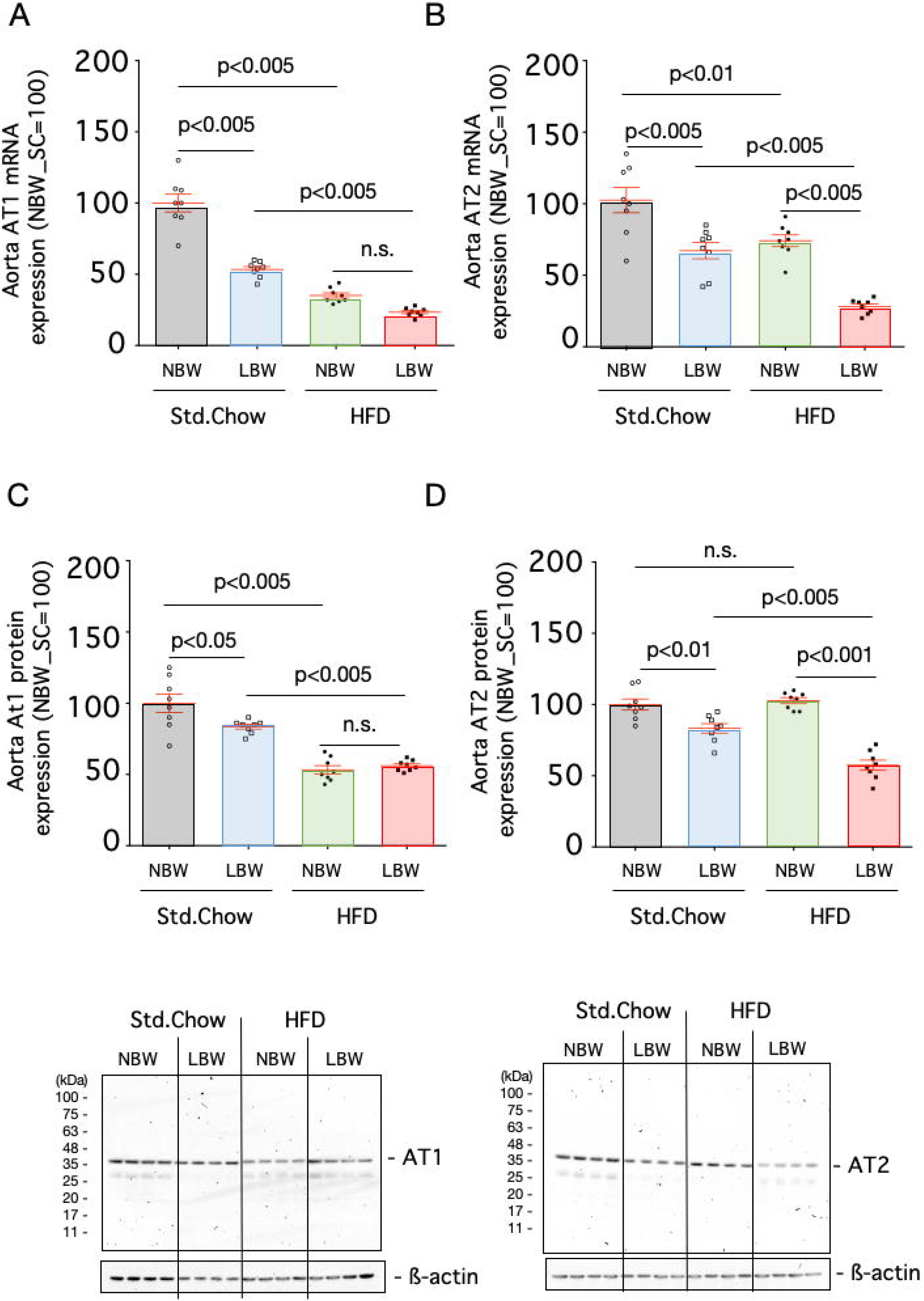
Expression of angiotensin receptor in the aorta. The mRNA expression levels of AT1 (A) and AT2 (B) and protein expression levels of AT1 (C) and AT2 (D) in the aortas of control rats (NBW) or LBW rats exposed to a standard chow or high-fat diet (HFD) were quantified. n=8.

**Figure 3.**
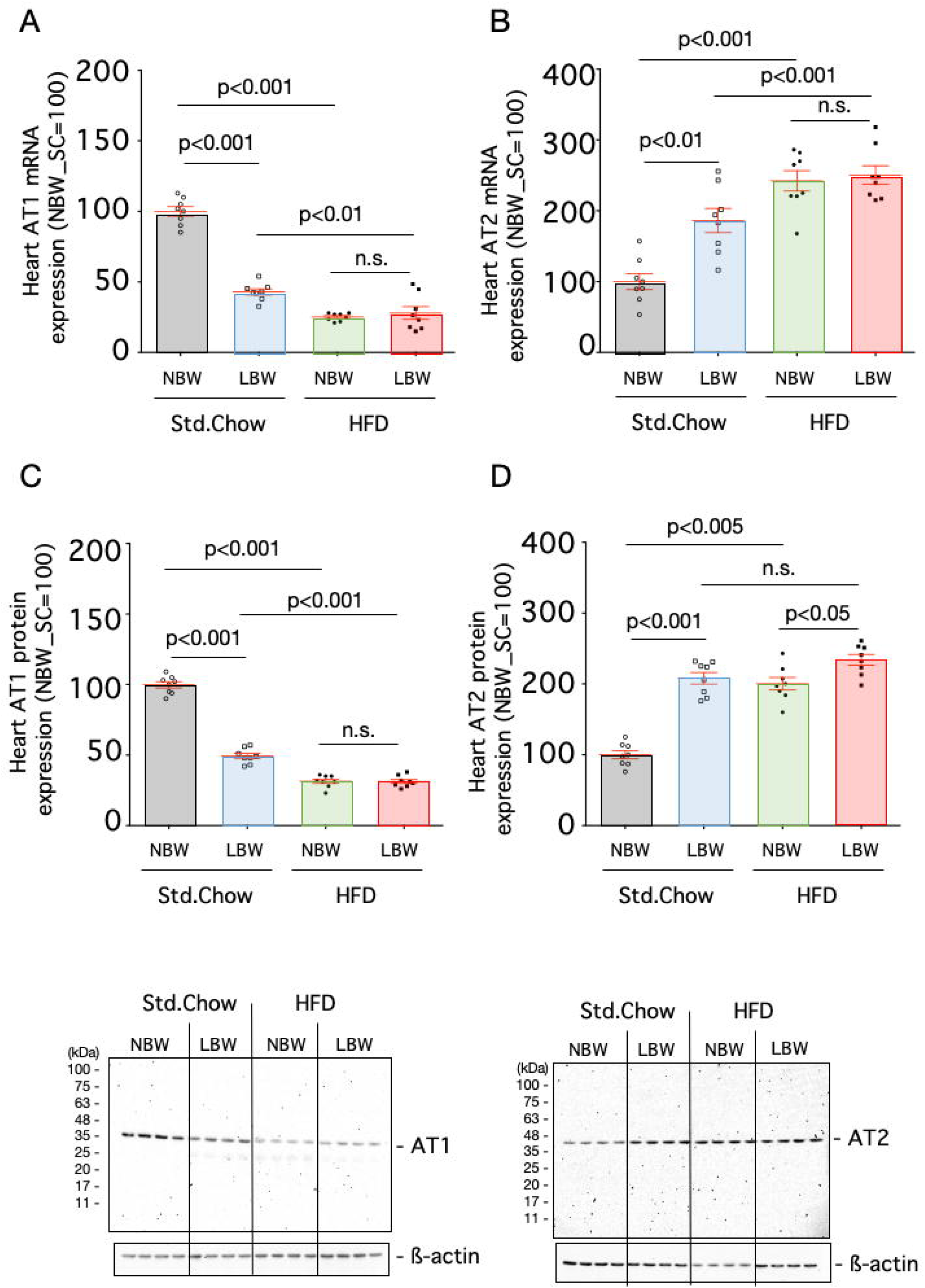
Expression of angiotensin receptor in the heart. The mRNA expression levels of AT1 (A) and AT2 (B) and protein expression levels of AT1 (C) and AT2 (D) in the hearts of control rats (NBW) or LBW rats exposed to a standard chow or high-fat diet (HFD) were quantified. n=8.

### Blood steroid levels

Blood aldosterone and corticosterone levels in LBW rats fed a standard diet did not differ from those of control rats (Fig. 4). When a high-fat diet was given to LBW rats, blood corticosterone levels increased significantly compared to control rats, but blood aldosterone levels did not differ between the two groups.

**Figure 4.**
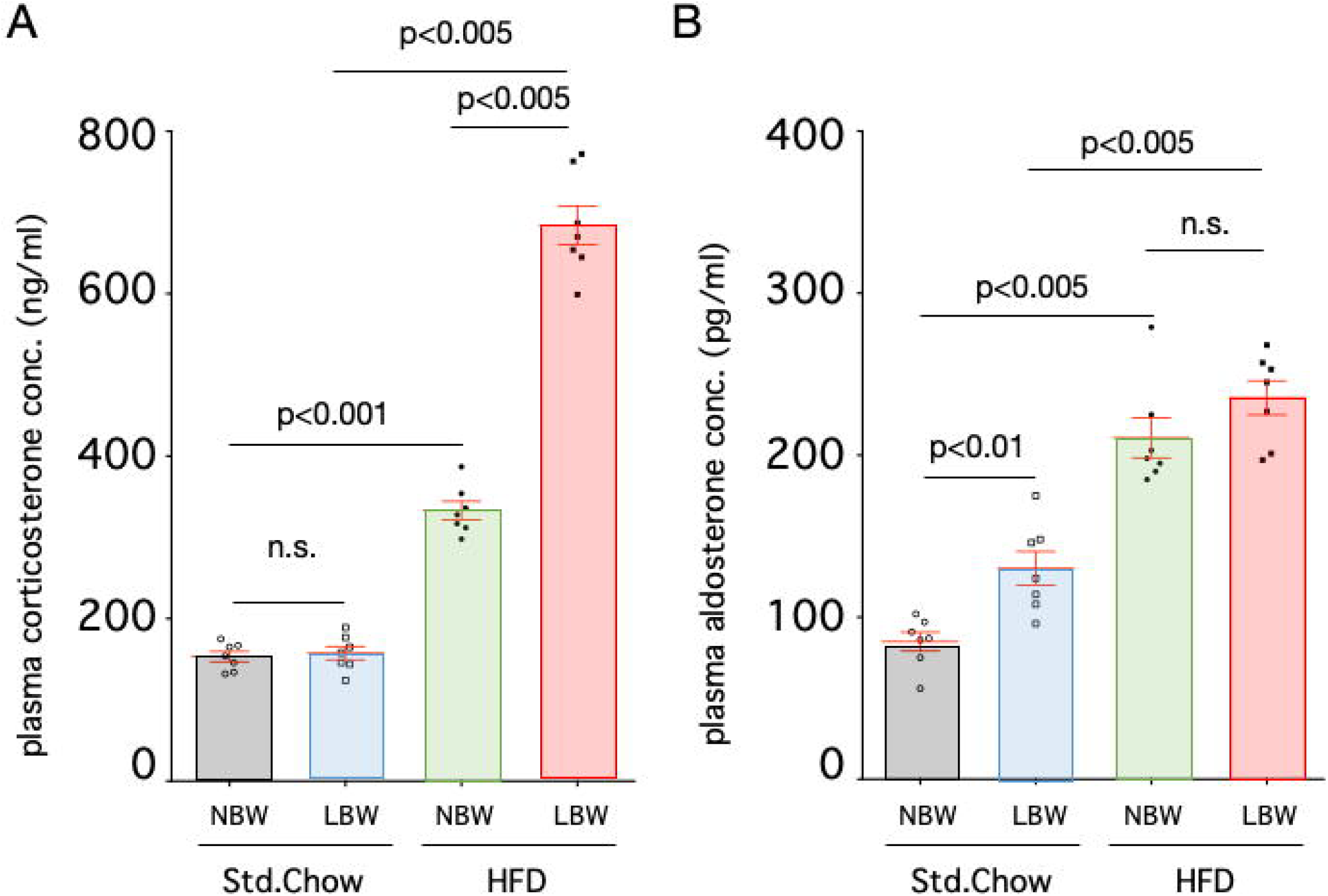
Blood aldosterone and corticosterone levels. Plasma concentrations of aldosterone (A) and corticosterone (B) of control rats (NBW) or LBW rats exposed to a standard chow or high-fat diet (HFD) were measured. n=8.

### Expression of miR-449a in the anterior pituitary

The expression of miR-449a in the pituitary glands of rats fed a standard diet was not different between control and LBW rats (Fig. 5). When a high-fat diet was given, pituitary miR-449a was increased compared to the standard diet in controls; however, in LBW rats there was no increase in miR-449a with high-fat diet exposure.

**Figure 5.**
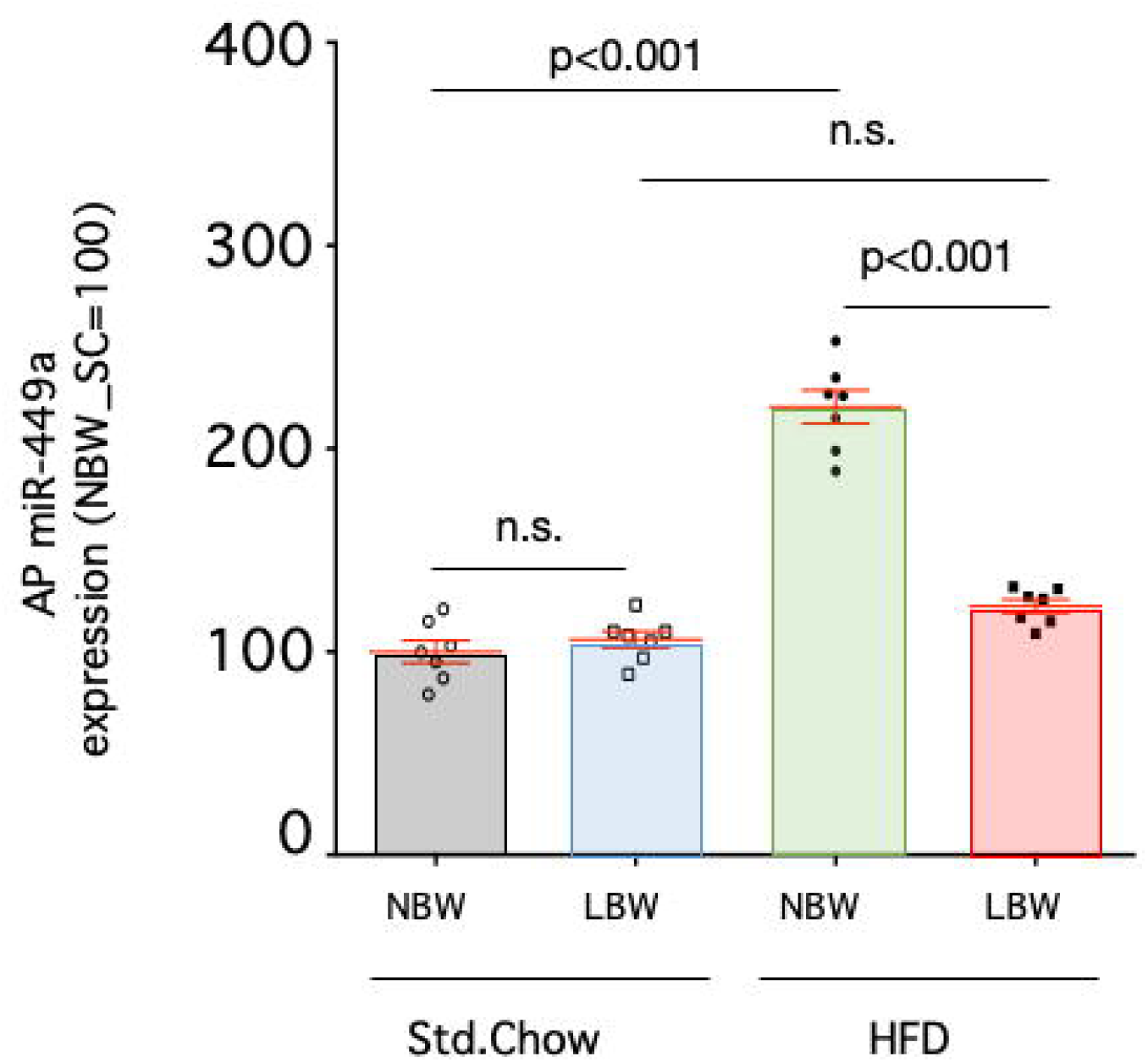
Expression of miR-449a in the anterior pituitary. Expression of miR-449a in the anterior pituitary of control rats (NBW) or LBW rats exposed to a standard chow or high-fat diet (HFD) were quantified. n=8.

### Administration of steroid synthesis inhibitors

Administration of metyrapone significantly reduced the upregulation of blood pressure in LBW rats to levels comparable with control rats (Fig. 6). Administration of metyrapone to control rats did not produce different results from vehicle administration.

**Figure 6.**
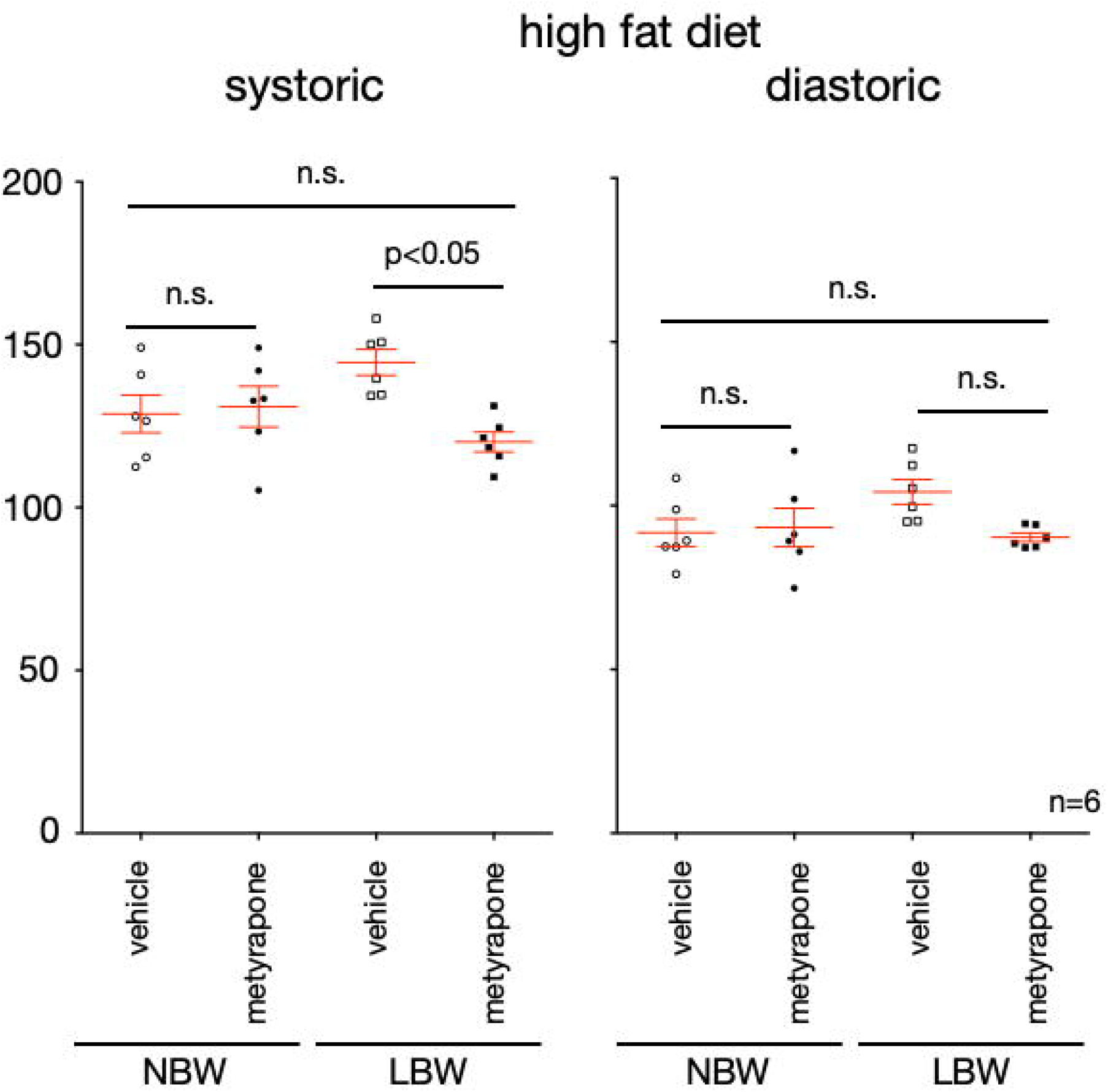
Changes in blood pressure after administration of steroid synthesis inhibitors. The blood pressure of control rats (NBW) and LBW rats exposed to the high-fat diet (HFD) after the administration of metyrapone or vehicle was measured by the tail cuff method. The diastolic blood pressure and the systolic blood pressure of each rat are shown. n=6.

## Discussion

In our rat model, elevated corticosterone levels following exposure to a high-fat diet may be responsible for increased blood pressure in LBW rats. In addition, the expression of AT1, which is involved in increasing blood pressure, also decreased. In contrast, AT2 expression, which is involved in decreasing blood pressure, was increased, and its alteration pattern is considered to be a compensatory change in the heart. All results, including the absence of pathological findings in the kidney and aorta as well as the elevation of blood aldosterone levels, indicated that renal and vascular diseases may not play a causative role in the development of hypertension in fetal low-carbohydrate and calorie-restricted rats.

It has been reported that premature babies have low nephron numbers because the formation of renal glomeruli depends on the embryonic period. In our model, there was no difference in gestational age, as previously reported^40^, and only birthweight was decreased. In fact, there was no difference in the weights of kidneys per body weight, and histological analysis of the kidney showed no differences in the number of glomeruli per visual field or in the morphological appearance. These findings indicate that the mechanism is different from the development of hypertension often seen in preterm infants. Furthermore, since the expression of AT1 was decreased in the hearts and aortas of high-fat diet-exposed LBW rats, these are considered to be compensatory changes. Although aldosterone levels in the blood increased with a high-fat diet compared to the standard chow, there was no difference between LBW and control rats. Moreover, there was no difference in heart rate. Although we did not investigate the functions of the autonomic nervous system, we consider that the involvement of the sympathetic nervous system may not be taken into account.

Protein intake is the key to increased blood pressure caused by fetal malnutrition. Maternal protein restriction during pregnancy has been reported to result in kidney abnormalities (decreased nephron number and glomerular morphology) and an increased renin-angiotensin system. Fujii *et al.* report that a 30% dietary restriction during pregnancy results in increased blood pressure in rat offspring, but branched-chain amino acid supplementation normalized it^46^. Those results show that the amino acids consumed by the mother are important for the formation of the fetal kidney. However, in our results, these abnormalities were not observed in our fetal low-carbohydrate calorie-restriction model. Although detailed alteration of metabolites from low-carbohydrate and calorie-restriction exposure needs to be investigated in the near future, we have shown that the mechanisms of hypertension are different between protein-restriction and carbohydrate-restriction.

Atherosclerosis has a long preclinical stage in which pathological changes appear in the arteries of children and young adults before overt clinical manifestations of the disease appear^47^. Both fetal and childhood nutrition is important in this process and has been shown to affect the risk of lifelong CVD (e.g. breastfeeding is associated with long-term cardiovascular risk factors). It has been reported that fetal high-density lipoprotein cholesterol (HDL-C) and total cholesterol (TC) levels are lower in IUGR, and that the atherogenic index is increased in IUGR compared to healthy neonates^48^. Arteriosclerosis is an inflammatory disease, and its onset is known to involve excessive LDL-C^49^ but may also involve a decrease in HDL-C. Because of the anti-inflammatory properties of HDL-C, a decrease in HDL-C during fetal development and an increase in atherogenic index may be involved in the future development of arteriosclerosis in IUGR children. Although we have not examined the blood lipid profile of this rat model, we did not observe thickening of vascular smooth muscle in the aorta. Furthermore, we found that the expression of AT2 was increased in the hearts of high-fat diet-exposed LBW rats compared to controls (Fig. 3B and D), but not in the aortas (Fig. 2B and D). AT2 is involved in the regulation of remodeling^30^. In the present study, the aortas of high-fat diet-exposed LBW rats may not have been damaged. However, future studies should examine whether the hearts of LBW rats exposed to a high-fat diet have some damage and what is involved in the mechanism of increased AT2 expression in the heart. In fact, there is a report indicating that a major risk factor for the development of arteriosclerosis as well as hypertension is preterm birth, and the risk of that in full-term small for gestational age (SGA) children is not high^50^. Therefore, it is unlikely that atherosclerosis is not a principal cause for hypertension observed in our model.

Excess glucocorticoids often survive inactivation by 11β-HSD-2 in the kidney and bind to mineralocorticoid receptors. As a result, blood pressure increases due to sodium reabsorption and increased fluid volume. Ingestion of a high-fat diet or obesity results in metabolic stress, leading to an increase in blood glucocorticoids^51, 52^. Excessive and prolonged glucocorticoid exposure is associated with a high cardiovascular and metabolic burden. Chronic hypercortisolemia causes more persistent visceral fat accumulation than high-fat diet-induced obesity^53^. Thus, prolonged exposure to high-fat diets can generate a negative loop of metabolic disease. We have previously reported that LBW rats show long-term high blood corticosterone after restraint stress, and that CRF-R1 fails to downregulate without inducing miR-449a expression in the pituitary gland^17, 18^. Since pituitary miR-449a is induced by glucocorticoids and downregulates corticotropin releasing factor receptor (CRF-R1) expression, miR-449a is thought to be one of the mechanisms of negative feedback of the HPA-axis by glucocorticoids. In the present study, we showed that blood corticosterone levels in high-fat diet-exposed LBW rats were higher than that in control rats. Pituitary miR-449a is induced in high-fat diet-exposed control rats compared to standard diet-fed control rats, but miR-449a did not increase in high fat diet-exposed LBW rats, and the mechanism is unknown. In addition, we have shown that administration of steroid synthesis inhibitors to high-fat diet-exposed LBW rats normalized their blood pressure. These results suggest that LBW rats may have abnormal pituitary glucocorticoid feedback of the HPA-axis due to metabolic stress induced by high-fat diet exposure, and blood pressure may have increased due to the increased blood corticosterone concentration.

In conclusion, we found that LBW rats had elevated blood pressure by high-fat diet exposure after growth. There are various causes of the risk of developing hypertension due to LBW, but it is necessary to develop a treatment strategy for hypertension for each cause of LBW. One of the mechanisms of hypertension in high-fat diet-exposed LBW is suggested to be related to elevated blood corticosterone levels in our model rats. We think that the mismatch between the acquired constitution by undernutrition and the environment induced by a high-fat diet after growth increases the risk of developing metabolic disease. Our results imply that LBW children resulting from low-carbohydrate and calorie-restricted diets in mothers should be carefully followed in terms of evaluating their blood cortisol levels. People suffering mild essential hypertension with a history of light weight at birth may need to reduce their daily fat intake as well as their salt intake.

## Author contributions

T. Nemoto designed the work, acquired data, analyzed data, and drafted the manuscript. T. Nakakura performed tissue analyses of kidneys and aortas. Y. Kakinuma interpreted the data and substantively revised the manuscript.

## Disclosure statement

The authors have nothing to disclose.

## Role of funding sources

This study was supported in part by JSPS KAKENHI Grant Number 17K10195.

## References

1. Lewington S, Clarke R, Qizilbash N, Peto R, Collins R, Prospective Studies C. Age-specific relevance of usual blood pressure to vascular mortality: a meta-analysis of individual data for one million adults in 61 prospective studies. Lancet. 2002;360(9349), 1903–1913.

2. Steinthorsdottir SD, Eliasdottir SB, Indridason OS, Palsson R, Edvardsson VO. The relationship between birth weight and blood pressure in childhood: a population-based study. Am J Hypertens. 2013;26(1), 76–82.

3. Huxley RR, Shiell AW, Law CM. The role of size at birth and postnatal catch-up growth in determining systolic blood pressure: a systematic review of the literature. J Hypertens. 2000;18(7), 815–831.

4. Barker DJ, Winter PD, Osmond C, Margetts B, Simmonds SJ. Weight in infancy and death from ischaemic heart disease. Lancet. 1989;2(8663), 577–580.

5. Paneth N, Susser M. Early origin of coronary heart disease (the “Barker hypothesis"). BMJ. 1995;310(6977), 411–412.

6. Klebanoff MA, Secher NJ, Mednick BR, Schulsinger C. Maternal size at birth and the development of hypertension during pregnancy: a test of the Barker hypothesis. Arch Intern Med. 1999;159(14), 1607–1612.

7. Katsuragi S, Okamura T, Kokubo Y, Ikeda T, Miyamoto Y. Birthweight and cardiovascular risk factors in a Japanese general population. J Obstet Gynaecol Res. 2017;43(6), 1001–1007.

8. Hoy WE, Nicol JL. The Barker hypothesis confirmed: association of LBW with all-cause natural deaths in young adult life in a remote Australian Aboriginal community. J Dev Orig Health Dis. 2019;10(1), 55–62.

9. Hales CN, Barker DJ. The thrifty phenotype hypothesis. Br Med Bull. 2001;60, 5–20.

10. Gluckman PD, Hanson MA, Beedle AS. Early life events and their consequences for later disease: a life history and evolutionary perspective. Am J Hum Biol. 2007;19(1), 1–19.

11. Gluckman PD, Hanson MA, Buklijas T. A conceptual framework for the developmental origins of health and disease. J Dev Orig Health Dis. 2010;1(1), 6–18.

12. Juvet C, Simeoni U, Yzydorczyk C, et al. Effect of early postnatal nutrition on chronic kidney disease and arterial hypertension in adulthood: a narrative review. J Dev Orig Health Dis. 2018;9(6), 598–614.

13. Hanson M. The Inheritance of Cardiovascular Disease Risk. Acta Paediatr. 2019; doi: 10.1111/apa.14813.

14. Kikuchi T, Uchiyama M. Epidemiological studies of the developmental origins of adult health and disease in Japan: a pediatric perspective in present day Japan. Clin Pediatr Endocrinol. 2010;19(4), 83–90.

15. Morisaki N, Urayama KY, Yoshii K, Subramanian SV, Yokoya S. Ecological analysis of secular trends in LBW births and adult height in Japan. J Epidemiol Community Health. 2017;71(10), 1014–1018.

16. Normile D. Staying slim during pregnancy carries a price. Science. 2018;361(6401), 440.

17. Nemoto T, Kakinuma Y, Shibasaki T. Impaired miR449a-induced downregulation of Crhr1 expression in LBW rats. J Endocrinol. 2015;224(2), 195–203.

18. Nemoto T, Kakinuma Y. Involvement of Noncoding RNAs in Stress-Related Neuropsychiatric Diseases Caused by DOHaD Theory: ncRNAs and DOHaD-Induced Neuropsychiatric Diseases. Adv Exp Med Biol. 2018;1012, 49–59.

19. Nuyt AM. Mechanisms underlying developmental programming of elevated blood pressure and vascular dysfunction: evidence from human studies and experimental animal models. Clin Sci (Lond). 2008;114(1), 1–17.

20. Nuyt AM, Alexander BT. Developmental programming and hypertension. Curr Opin Nephrol Hypertens. 2009;18(2), 144–152.

21. Ligi I, Grandvuillemin I, Andres V, Dignat-George F, Simeoni U. LBW infants and the developmental programming of hypertension: a focus on vascular factors. Semin Perinatol. 2010;34(3), 188–192.

22. Luyckx VA, Bertram JF, Brenner BM, et al. Effect of fetal and child health on kidney development and long-term risk of hypertension and kidney disease. The Lancet. 2013;382(9888), 273–283.

23. Kambayashi Y, Bardhan S, Takahashi K, et al. Molecular cloning of a novel angiotensin II receptor isoform involved in phosphotyrosine phosphatase inhibition. J Biol Chem. 1993;268(33), 24543–24546.

24. Mukoyama M, Nakajima M, Horiuchi M, Sasamura H, Pratt RE, Dzau VJ. Expression cloning of type 2 angiotensin II receptor reveals a unique class of seven-transmembrane receptors. J Biol Chem. 1993;268(33), 24539–24542.

25. Wang ZQ, Moore AF, Ozono R, Siragy HM, Carey RM. Immunolocalization of subtype 2 angiotensin II (AT2) receptor protein in rat heart. Hypertension. 1998;32(1), 78–83.

26. Miyata N, Park F, Li XF, Cowley AW, Jr. Distribution of angiotensin AT1 and AT2 receptor subtypes in the rat kidney. Am J Physiol. 1999;277(3), F437–446.

27. Kumar V, Knowle D, Gavini N, Pulakat L. Identification of the region of AT2 receptor needed for inhibition of the AT1 receptor-mediated inositol 1,4,5-triphosphate generation. FEBS Lett. 2002;532(3), 379–386.

28. Widdop RE, Jones ES, Hannan RE, Gaspari TA. Angiotensin AT2 receptors: cardiovascular hope or hype? Br J Pharmacol. 2003;140(5), 809–824.

29. Jones ES, Vinh A, McCarthy CA, Gaspari TA, Widdop RE. AT2 receptors: functional relevance in cardiovascular disease. Pharmacol Ther. 2008;120(3), 292–316.

30. Ludwig M, Steinhoff G, Li J. The regenerative potential of angiotensin AT2 receptor in cardiac repair. Can J Physiol Pharmacol. 2012;90(3), 287–293.

31. Xu J, Sun Y, Carretero OA, et al. Effects of cardiac overexpression of the angiotensin II type 2 receptor on remodeling and dysfunction in mice post-myocardial infarction. Hypertension. 2014;63(6), 1251–1259.

32. Masaki H, Kurihara T, Yamaki A, et al. Cardiac-specific overexpression of angiotensin II AT2 receptor causes attenuated response to AT1 receptor-mediated pressor and chronotropic effects. J Clin Invest. 1998;101(3), 527–535.

33. Akishita M, Horiuchi M, Yamada H, et al. Inflammation influences vascular remodeling through AT2 receptor expression and signaling. Physiol Genomics. 2000;2(1), 13–20.

34. Li J, Culman J, Hortnagl H, et al. Angiotensin AT2 receptor protects against cerebral ischemia-induced neuronal injury. FASEB J. 2005;19(6), 617–619.

35. Altarche-Xifro W, Curato C, Kaschina E, et al. Cardiac c-kit+AT2+ cell population is increased in response to ischemic injury and supports cardiomyocyte performance. Stem Cells. 2009;27(10), 2488–2497.

36. Curato C, Slavic S, Dong J, et al. Identification of noncytotoxic and IL-10-producing CD8+AT2R+ T cell population in response to ischemic heart injury. J Immunol. 2010;185(10), 6286–6293.

37. Visentin S, Grumolato F, Nardelli GB, Di Camillo B, Grisan E, Cosmi E. Early origins of adult disease: LBW and vascular remodeling. Atherosclerosis. 2014;237(2), 391–399.

38. Nemoto T, Mano A, Shibasaki T. miR-449a contributes to glucocorticoid-induced CRF-R1 downregulation in the pituitary during stress. Mol Endocrinol. 2013;27(10), 1593–1602.

39. Nemoto T, Mano A, Shibasaki T. Increased expression of miR-325-3p by urocortin 2 and its involvement in stress-induced suppression of LH secretion in rat pituitary. Am J Physiol Endocrinol Metab. 2012;302(7), E781–787.

40. Nemoto T, Kakinuma Y. Fetal malnutrition-induced catch up failure is caused by elevated levels of miR-322 in rats. Sci Rep. 2020;10(1), 1339.

41. Nakakura T, Suzuki T, Nemoto T, et al. Intracellular localization of alpha-tubulin acetyltransferase ATAT1 in rat ciliated cells. Med Mol Morphol. 2016;49(3), 133–143.

42. Wang Y, Thatcher SE, Cassis LA. Measuring Blood Pressure Using a Noninvasive Tail Cuff Method in Mice. Methods Mol Biol. 2017;1614, 69–73.

43. Aoyagi Y, Furuyama T, Inoue K, et al. Attenuation of Angiotensin II-Induced Hypertension in BubR1 Low-Expression Mice Via Repression of Angiotensin II Receptor 1 Overexpression. J Am Heart Assoc. 2019;8(23), e011911.

44. Nolan T, Hands RE, Bustin SA. Quantification of mRNA using real-time RT-PCR. Nat Protoc. 2006;1(3), 1559–1582.

45. Parthasarathy C, Yuvaraj S, Sivakumar R, Ravi Sankar B, Balasubramanian K. Metyrapone-induced corticosterone deficiency impairs glucose oxidation and steroidogenesis in Leydig cells of adult albino rats. Endocr J. 2002;49(4), 405–412.

46. Fujii T, Yura S, Tatsumi K, et al. Branched-chain amino acid supplemented diet during maternal food restriction prevents developmental hypertension in adult rat offspring. J Dev Orig Health Dis. 2011;2(3), 176–183.

47. Singhal A. The early origins of atherosclerosis. Adv Exp Med Biol. 2009;646, 51–58.

48. Pecks U, Brieger M, Schiessl B, et al. Maternal and fetal cord blood lipids in intrauterine growth restriction. J Perinat Med. 2012;40(3), 287–296.

49. Ross R. Atherosclerosis--an inflammatory disease. N Engl J Med. 1999;340(2), 115–126.

50. Rossi P, Tauzin L, Marchand E, Boussuges A, Gaudart J, Frances Y. Respective roles of preterm birth and fetal growth restriction in blood pressure and arterial stiffness in adolescence. J Adolesc Health. 2011;48(5), 520–522.

51. Khazen T, Hatoum OA, Ferreira G, Maroun M. Acute exposure to a high-fat diet in juvenile male rats disrupts hippocampal-dependent memory and plasticity through glucocorticoids. Sci Rep. 2019;9(1), 12270.

52. Puigoriol-Illamola D, Leiva R, Vazquez-Carrera M, Vazquez S, Grinan-Ferre C, Pallas M. 11beta-HSD1 Inhibition Rescues SAMP8 Cognitive Impairment Induced by Metabolic Stress. Mol Neurobiol. 2019; doi: 10.1007/s12035-019-01708-4.

53. Garcia-Eguren G, Giro O, Romero MDM, Grasa M, Hanzu FA. Chronic hypercortisolism causes more persistent visceral adiposity than HFD-induced obesity. J Endocrinol. 2019;242(2), 65–77.

